# Hypothalamic tau^P301L^ pathology disrupts pair bonding in monogamous prairie voles

**DOI:** 10.64898/2026.09.28.755178

**Authors:** Michael A. Kelberman, Kelly E. Winther, Razan E. Ibrahiem, Eric G. Albregts, Meenakshi Pardasani, Yurika L. Watanabe, Zoe R. Donaldson

## Abstract

Although dementias primarily affect memory, they also disrupt our closest social relationships, resulting in elevated caregiver burden and worse patient outcomes. However, the mechanistic study of how pathology affects social attachments is limited by the use of laboratory rodents that do not form selective, long-term bonds. Here, in monogamous prairie voles, we expressed dementia-causing tau^P301L^ in the paraventricular hypothalamus of 12- and 18-month-old animals, and separately, from birth through 18-months, across hippocampocortical regions. Hippocampocortical expression of tau^P301L^ did not yield deficits in social or non-social dementia-related phenotypes. In contrast, tau^P301L^ expressed in the hypothalamus yielded sex- and age-dependent social bonding deficits. 18-month males with hypothalamic tau pathology failed to form bonds and displayed blunted sociability, effects that converged across bonding assays. In addition, healthy 18-month males paired with a hypothalamic tau-expressing female also showed reduced partner preference, suggesting that pathology exerts vicarious effects on partners. Social behavior of both healthy females and those expressing tau^P301L^ were unaffected. 18-month tau^P301L^ males also showed elevated oxytocin fluorescence. These findings establish direct and vicarious disruptions to pair bonding induced by hypothalamic tau pathology in male prairie voles, potentially via altered oxytocin signaling.

## Introduction

Dementias are a diverse collection of diseases traditionally characterized by neuronal loss and cognitive impairment, yet dementia patients often develop a constellation of non-cognitive symptoms particularly during the prodromal phases of disease^1^. Among the most prevalent and persistent of these non-cognitive symptoms are those that strain social relationships, including social withdrawal, irritability, mood changes, and agitation^1–4^. These symptoms are associated with poorer outcomes, greater functional impairment, and accelerated decline for patients, and also place the heavier burden on social support systems^1,5–7^.

The non-cognitive behavioral impairments that occur in dementia patients make caregiving particularly challenging, especially when the caregiver is also the spouse whose relationship norms can be shattered by the changes in their partner. Spouses frequently provide more caregiving hours than adult children, experience the highest levels of caregiver burden, and face an elevated risk of developing dementia themselves when caring for a spouse with the disease^8–15^. Moreover, dementia-induced reductions in relationship quality emerge at a pivotal point in disease course. The same social bonds strained by dementia are among the most potent modifiers of disease progression: strong social support can preserve cognition and slow disease progression even in the face of dementia-causing pathology^16–19^, while events such as the death of a spouse can promote cognitive decline and dementia beyond the deleterious impacts of general loneliness and social isolation^12,20–22^. Dementia therefore unfolds within the context of social bonds, ultimately placing spousal relationships at the center of both risk and resilience.

Dementia symptoms have traditionally been tied to hippocampal and cortical dysfunction^23–27^, consistent with the classical model of pathology in these regions driving cell loss and cognitive decline. However, non-cognitive behavioral deficits that typify dementia, including social symptoms, are often attributable to tau-induced dysfunction of subcortical structures^28–33^. Among the constellation of subcortical systems affected in dementias, the paraventricular hypothalamus (PVH) represents a potential nexus underlying dementia-related social deficits, specifically. Distinct PVH cell populations synthesize corticotropin releasing hormone, vasopressin, and oxytocin (OXT), among other neuropeptides. Many of these are implicated in social behavior, including the complex attachment behaviors that define spousal relationships^34–38^. The PVH also displays early vulnerability to accumulation of tau pathology in dementia^39–42^, though evidence linking PVH neural degeneration to symptoms is mixed^40,43–46^. Even so, treatments modulating PVH-derived neurochemicals have shown benefit across multiple rodent models of dementia, including improving social deficits^32,47–52^. Together, these observations suggest that tau-induced PVH dysfunction may contribute to broad social deficits in dementia, but its specific involvement in attachment-related deficits remains presumptive largely because existing preclinical models (e.g. mice and rats) do not form adult bonds and are insufficient for testing this.

Monogamous prairie voles, which form enduring pair bonds marked by partner preference and aggression toward unfamiliar conspecifics^38,53^, offer a promising alternative for asking both how PVH pathology drives attachment-related deficits and how it affects a bonded, pathology-free partner. We have previously shown age-dependent shifts in pair bonding: 18-month voles form pair bonds faster (males) and display increased partner directed affiliative behavior (females) than younger voles^54^, mirroring other aspects of behavior that are dependent on age^55–58^.

Humans similarly report that the quality of their social relationships improves with age, driven in part by increased positive emotions and satisfaction within those established relationships^59,60^. We also found that augmented bonding co-occurred with increased PVH OXT^+^, but not PVH vasopressin^+^, cell number in 18-month voles, suggesting that behavioral changes during aging are supported by the same PVH-derived neuromodulators that are potentially disrupted by dementia pathophysiology.

Here, we tested the hypothesis that PVH rather than hippocampocortical dementia-related tau pathology is a primary driver of attachment-related social deficits. We overexpressed tau^P301L^, a rare, autosomal dominant dementia-inducing mutation^61–64^ in these brain regions and examined social and non-social behaviors in aged voles. Tau^P301L^ expressed throughout hippocampal and cortical regions from birth did not yield deficits in any of a suite of social or non-social phenotypes. In contrast, tau^P301L^ expressed in the PVH yielded sex- and age-related deficits in pair bonding; PVH tau^P301L^ pathology impaired bond formation and reduced sociability in 18- month but not 12-month males. Males paired with PVH tau^P301L^ females also exhibited bonding deficits, suggesting pathology-driven changes in one partner produce vicarious effects in a healthy partner. In contrast, females were largely resilient to the direct and vicarious effects of PVH tau^P301L^ pathology on partner preference and sociability. Non-social behaviors were largely unaffected in both sexes. Although tau^P301L^ was mainly expressed in non-OXT neurons, we identified elevated OXT immuno- reactivity in 18-month PVH males, suggesting potential indirect oxytocinergic effects of PVH tau^P301L^ expression. These findings directly implicate hypothalamic tau pathology in sex- and age- dependent bond disruption that impacts both members of a pair via mechanisms that may involve altered OXT signaling.

## Results

### Region-specific viral expression of tau^P301L^ produces restricted pathology

We developed viral strategies to differentiate the effects of tau^P301L^ pathology in hippocampocortical regions versus the PVH specifically. Viral injections into the transverse sinus on postnatal day 1 (P1/neonate) resulted in the expression of phosphorylated tau in cortical and hippocampal regions while sparing the PVH (**Fig. 1C**; **Fig. S1A-F**)^65^, and intra-PVH infusions produced local expression of phosphorylated tau in both 12- and 18-month animals (**Fig. 1C**; **Fig. S1G,H**). Voles with hippocampocortical targeting did not exhibit phosphorylated tau levels beyond that observed in control animals (**Fig. 1D**). In contrast, 12-month PVH tau^P301L^ animals exhibited more phosphorylated tau expression within the PVH (One-way ANOVA, main effect of treatment: Males: F_2,28_ = 3.894, p = 0.0322; Females: F_2,28_ = 4.200, p = 0.0254; **Fig. 1E**)

**Fig. 1:**
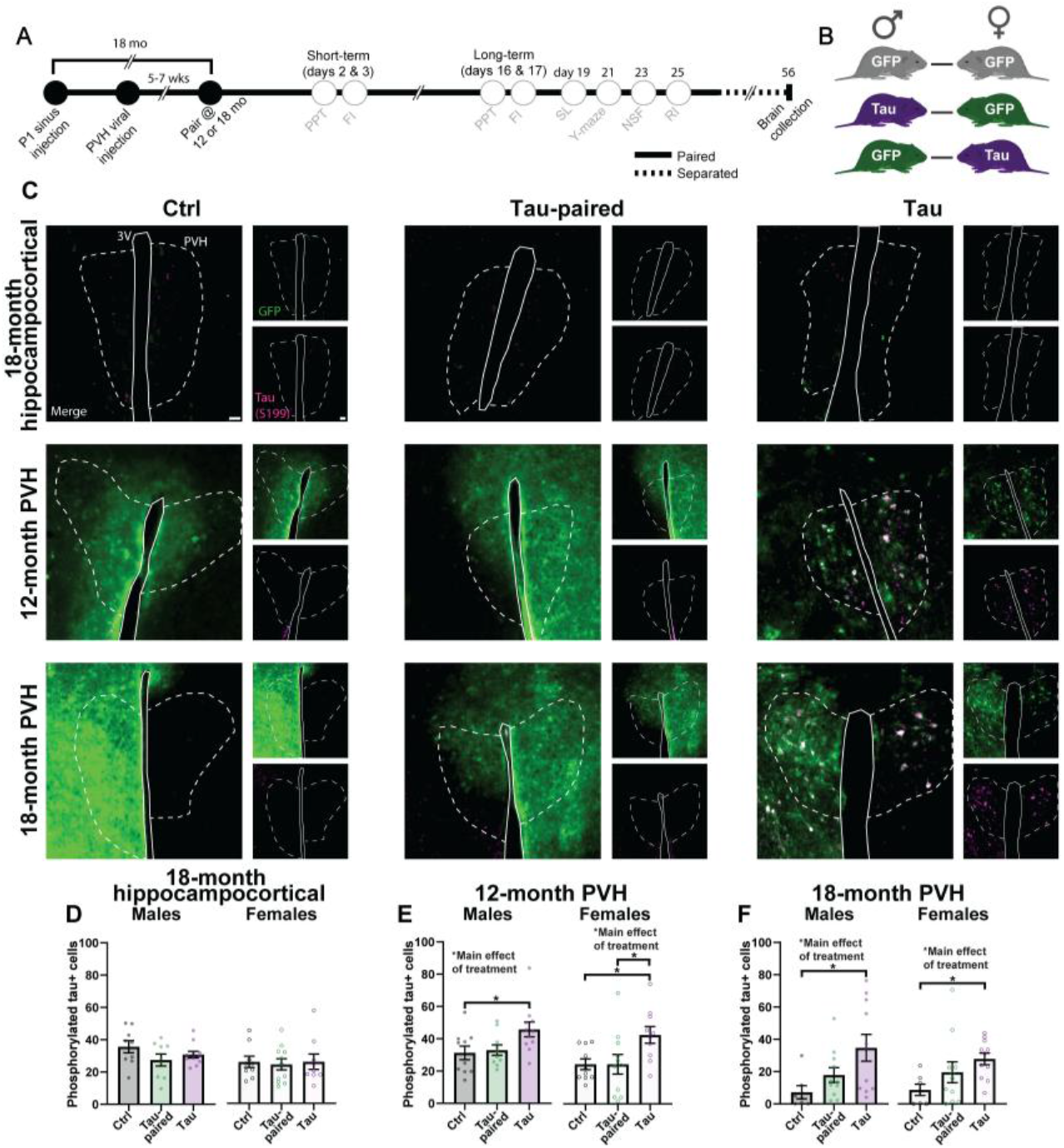
Viral strategies produce two distinct expression patterns of tau^P301L^ pathology. **A** Experimental timeline. PVH: paraventricular hypothalamus, PPT: partner preference test, FI: free interaction, SL: sleep latency, NSF: novelty suppressed feeding, RI: resident intruder. **B** Pairing scheme. **C** Representative images of GFP (green) and phosphorylated tau (S199, magenta) staining in the PVH from hippocampocortical and PVH targeted animals across the three pairing types (ctrl, tau-paired, and tau). Scale = 50 µm. **D-E** Hippocampocortical injections did not result in appreciable PVH phosphorylated tau whereas targeted PVH injections produced elevated phosphorylated tau in males and females at 12- and 18-months. *p<0.05.

which manifested as males accumulating more phosphorylated tau than control animals (Tukey’s post-hoc: q_28_ = 3.581, p = 0.0441) and females accumulating more than both control (Tukey’s post-hoc: q_28_ = 3.521, p = 0.0484) and tau-paired (Tukey’s post-hoc: q_28_ = 3.604, p = 0.0425) counterparts. PVH tau^P301L^ expression in 18-month animals also led to greater phosphorylated tau levels (Kruskal-Wallis test, main effect of treatment: Males: Kruskal-Wallis statistic_3,29_ = 8.063, p = 0.0177; Females: Kruskal-Wallis statistic_3,29_ = 7.488, p = 0.0237; **Fig. 1F**). For both males and females, there was a graded effect with the lowest tau detected in controls, an intermediate amount in tau-paired animals, and the greatest levels in tau_P301L_ animals. Pairwise comparisons showed that 18-month PVH tau^P301L^ males and females accumulated more tau compared to control animals (Dunn’s post-hoc: Males: Z_7,11_ = 2.840, p = 0.0136; Females: Z_7,11_ = 2.673, p = 0.0225); tau-paired voles did not differ from either control or tau_P301L_ animals (Dunn’s post-hoc: Males: Z_11,11_ = 1.252, p = 0.6316; Females: Z_11,11_ = 1.704, p = 0.2654).

### Tau^P301L^ expression in male voles disrupts bonding in an age- and region-dependent manner

We next asked whether our viral manipulations resulted in differences in pair bonding (**Fig. 2**; **Fig. S2**). Hippocampocortical tau^P301L^ tau expression had no significant effects on partner preference metrics (partner preference index or partner, stranger, or total huddle time) in 18-month male voles (**Fig. 2C**; **Fig. S2C-E**). 12-month PVH tau^P301L^ male voles similarly did not exhibit any differences in partner preference metrics (**Fig. 2D**; **Fig. S2F-H**). However, PVH tau^P301L^ expression impaired partner preference in 18-month males (Two-way ANOVA, main effect of treatment: F_2,26_ = 3.665, p = 0.0396; **Fig. 2E**). While 18-month control males formed strong partner preferences, both tau-paired and tau^P301L^ males displayed lower partner preference regardless of timepoint, suggesting both a main and vicarious effect of tau pathology on attachment behaviors. To confirm a lack of partner preference, we followed up with a one-sample t-test comparing partner preference to chance (e.g. 0.5). 18-month PVH control males displayed strong partner preference at the short- (t_6_ = 13.30, p < 0.0001) and long-term (t_6_ = 92.40, p < 0.0001) timepoint. While 18-month PVH tau-paired males did not show short-term partner preference (t_10_ = 1.128, p = 0.2856), they did form a preference after long-term cohabitation (t_10_ = 3.059, p = 0.0121). Meanwhile, 18-month PVH tau^P301L^ animals did not form a partner preference at either timepoint (short-term: t_10_ = 1.569, p = 0.1476; long-term: t_10_ = 1.527, p = 0.1578). Reduced partner preference was not due to changes in the amount of time that tau-paired and tau^P301L^ animals spent huddling with their partner (**Fig. S2I**); rather both groups displayed elevated time huddling with the stranger animal (Two-way ANOVA, main effect of treatment: F_2,26_ = 6.358, p = 0.0057; **Fig. S2J**). Finally, 18-month tau^P301L^ males were less social, spending less total time huddling with either animal during the compared to control and tau-paired males (Two-way ANOVA, main effect of treatment: F_2,26_ = 4.914, p = 0.0155; **Fig S2K**).

**Fig. 2:**
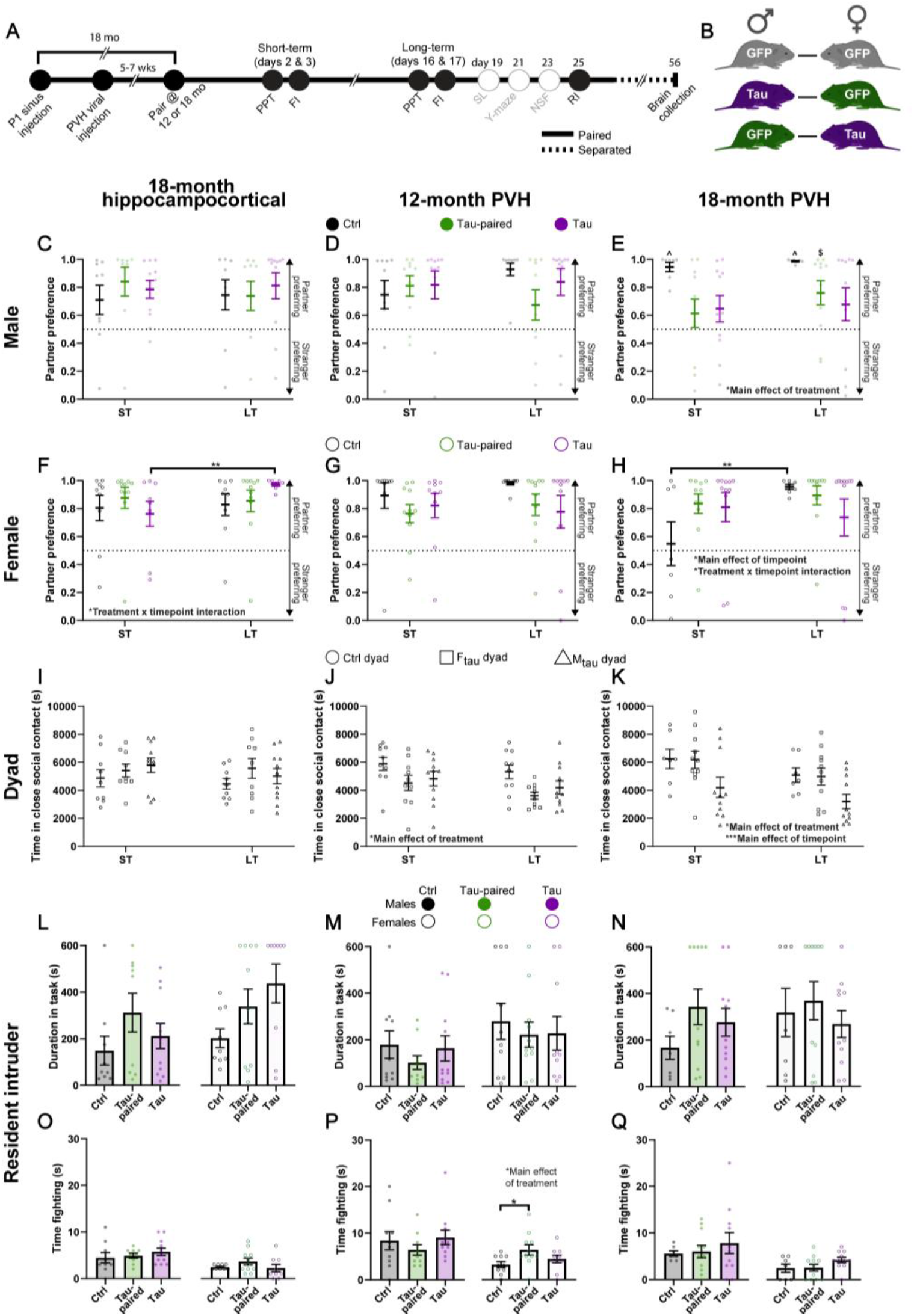
Tau^P301L^ pathology exerts direct and vicarious effects on pair bonding in a sex-, age-, and region-specific manner. **A** Experimental timeline. PVH: paraventricular hypothalamus, PPT: partner preference test, FI: free interaction, RI: resident intruder. **B** Pairing scheme. **C-E** Partner preference scores in 18-month hippocampocortical and 12- and 18-month PVH males. 18-month PVH tau-paired and tau^P301L^ males displayed lower partner preference compared to controls. Dashed line = chance (0.5). ^p<0.0001 compared to chance (0.5), ^$^p<0.05 compared to chance (0.5). **F-H** Partner preference scores in 18-month hippocampocortical and 12- and 18-month PVH females. 18-month tau^P301L^ hippocampocortical and 18-month PVH control females increased their partner preference from the short- to long-term timepoint. Dashed line = chance (0.5). **I-K** Time spent in close social contact during unconstrained dyadic interaction. 12-month PVH pairs with a tau^P301L^ animal, regardless of sex, displayed lower time spent in close social contact. 18-month PVH pairs displayed lower time spent in close social contact at the long-term timepoint and 18-month PVH pairs with a tau^P301L^ male displayed the lowest time spent in close social contact, regardless of timepoint. **L-Q** Aggression measured by the resident intruder task. Only 12-month PVH females displayed a main effect of treatment, with tau-paired animals spending longer fighting than controls. ST: short-term, LT: long-term; *p<0.05, **p<0.01, ***p<0.001.

### Female attachment behaviors are less sensitive to tau^P301L^ expression and pairing with a tau^P301L^ partner

Cohabitation length, rather than tau^P301L^ expression and pairing, was the primary driver of differences in partner preference in female voles. Specifically, 18-month females with hippocampocortical pathology displayed a timepoint x treatment interaction (Two-way ANOVA, timepoint x treatment interaction: F_2,26_ = 3.724, p = 0.0378; **Fig. 2F**) where tau^P301L^ females increased their partner preference from the short- to longterm timepoint (Holm-Šidák post-hoc: t_26_ = 3.207, p = 0.0106). The amount of time spent huddling with a partner generally increased with cohabitation length (Two-way ANOVA, main effect of timepoint: F_1,26_ = 4.230, p = 0.0499; **Fig. S3C**), which was most apparent in tau^P301L^ animals. No main or interacting effects were noted in either stranger or total huddle time in 18-month females with hippocampocortical expression (**Fig. S3D,E**).

There were no main or interacting effects in 12- month PVH females on any partner preference test metric (**Fig. 2G, Fig. S3F-H**), but in 18-month PVH females, there was a significant main effect of timepoint (Two-way ANOVA, main effect of treatment: F_1,26_ = 4.640, p = 0.0407) and a timepoint x treatment interaction (Two-way ANOVA, timepoint x treatment interaction: F_2,26_ = 4.903, p = 0.0156) on partner preference (**Fig. 2H**). Post-hoc testing revealed that 18-month PVH control females increased their partner preference from the short- to long-term timepoint (Holm-Šidák post-hoc: t_26_ = 3.360, p = 0.0072). There was also a main effect of timepoint on partner huddle time (Two-way ANOVA, main effect of timepoint: F_1,26_ = 6.408, p = 0.0178; **Fig. S3I**), where 18- month PVH females tended to increase the time spent huddling with their partners, which was most evident in control and tau-paired animals. No main effects were noted in either stranger or total huddle time in 18-month PVH females (**Fig. S3J,K**).

### Unconstrained social interaction reveals the influence of tau expression and tau-pairing on intra-pair behavioral patterns

Partner preference employs social choice as a proxy for bonding. However, this behavioral paradigm is not well suited to examine bond-related behaviors when both members of the pair have equal agency and opportunities to engage or disengage. Accordingly, the behaviors expressed during partner preference tests are only nominally associated with those observed during unconstrained dyadic interaction^54,66^. To examine the effects of tau^P301L^ on unconstrained pairwise behavior, we performed free interaction tests the day after each partner preference test (**Fig. 2I-K**), recording the total amount of time spent in close social contact across 3 hrs. We observed no effect of pathology on time spent in close social contact in pairs expressing hippocampocortical pathology (**Fig. 2I**). 12- month pairs that had a tau^P301L^ expressing animal, regardless of which sex expressed the pathology, spent less time in close social contact (Two-way ANOVA, main effect of treatment: F_2,26_ = 5.187, p = 0.0121; **Fig. 2J**). In 18-month PVH animals, pairs displayed decreased amount of time spent in close social contact after long-term cohabitation, and pairs with a male tau^P301L^ animal tended to show the lowest time spent in close social contact regardless of timepoint (Two-way ANOVA, main effect of timepoint: F_1,26_ = 16.43, p = 0.0004; Two-way ANOVA, main effect of treatment: F_2,26_ = 3.777, p = 0.0363; **Fig. 2K**).

### Resident intruder-elicited aggression was similar across treatment groups regardless of changes in bonding

Finally, we employed a resident intruder task as a readout of another facet of pair bonding (**Fig. 2L-Q**), as bonded voles display aggression towards a same-sex conspecific. All groups completed the task at similar rates (5 tumble fights, 30 s total fighting, or 10 min; **Fig. 2L-N**), and 12-month PVH females were the only group to display changes in the amount of time spent fighting (One-way ANOVA, main effect of treatment: F_2,28_ = 3.357, p = 0.0493; **Fig. 2O-Q**), tau-paired animals spent more time fighting than controls (Tukey’s post-hoc: q_28_ = 3.616, p = 0.0418).

### Tau^P301L^ expression does not affect other, non-social dementia-related phenotypes

To determine whether tau-induced deficits were most evident in the social domain, we employed a suite of behavioral assays aimed at assessing non-social deficits common in dementia (**Fig. 3**). Overall, non- social behavioral phenotypes were unaffected by either hippocampocortical or PHV tau^P301L^ pathology.

**Fig. 3:**
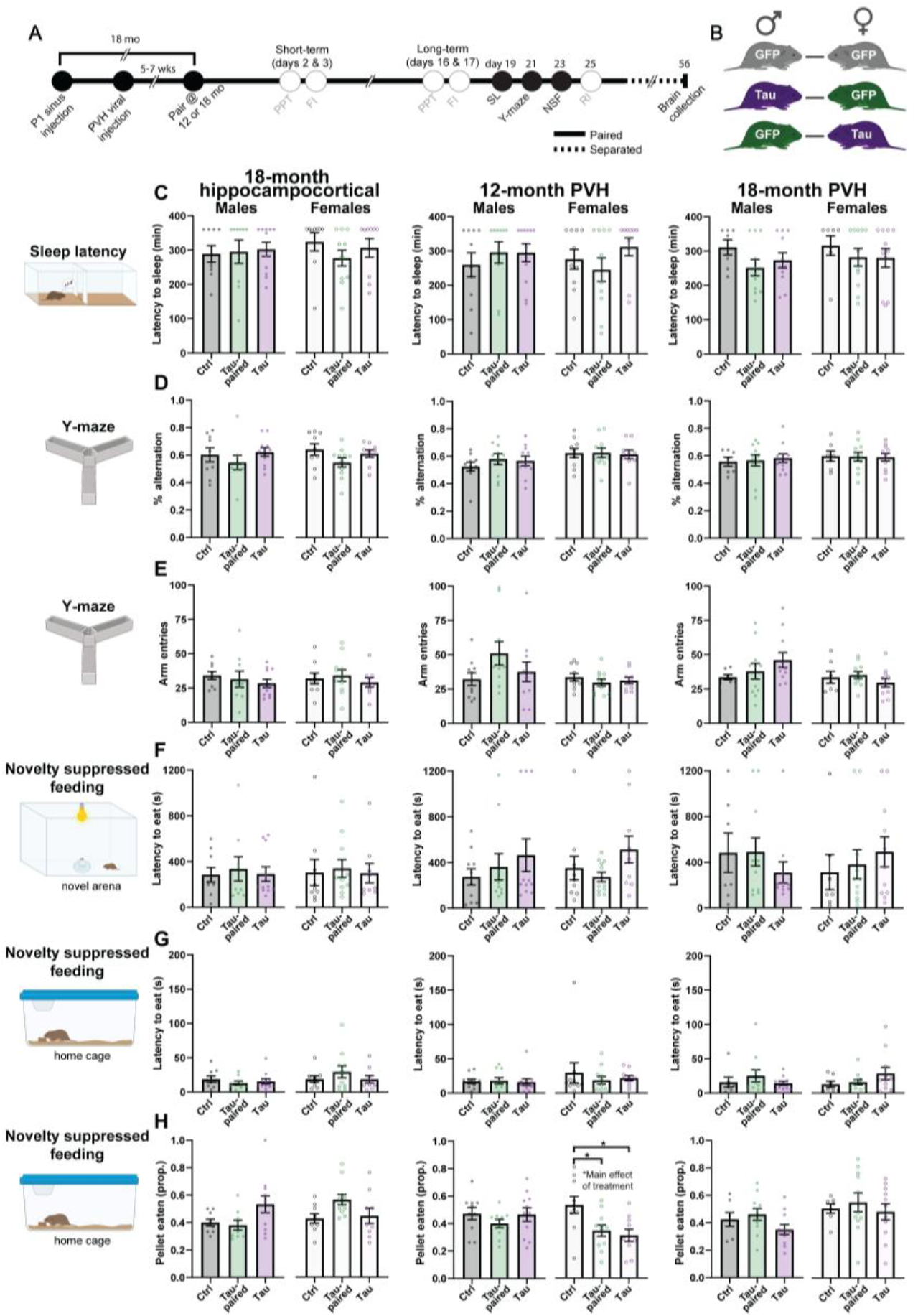
Tau^P301L^ pathology has minimal effect on non-social behaviors. **A** Experimental timeline. SL: sleep latency, NSF: novelty suppressed feeding. **B** Pairing scheme. **C** Latency to sleep measured during a sleep latency task revealed no effects of treatment across any group. **D-E** Working memory (D) and locomotor activity (E) measured during a Y-maze task revealed no effects of treatment across any group. **F-H** Latency to eat in a novel environment (F) and home cage (G), and amount eaten in a 30-min span following novelty suppressed feeding (H). 12-month PVH females displayed a main effect of treatment, with controls eating more than tau-paired and tau^P301L^ animals in the 30-min period following novelty suppressed feeding. *p<0.05.

Briefly, neither general arousal (sleep latency) nor spatial memory (y-maze) were affected by any of our treatments (**Fig. 3C-E**). Anxiety-like behavior (novelty suppressed feeding) did not yield any differences in the latency to eat in a novel environment or the home cage, and all groups consumed the same amount of food within a 30-min period following the task (**Fig. 3F-H**). The one exception was in 12-month PVH females (One-way ANOVA, main effect of treatment: F_2,28_ = 5.699, p = 0.0084), where control animals ate more than both tau-paired (Tukey’s post-hoc: q_28_ = 3.815, p = 0.0305) and tau^P301L^ (Tukey’s post-hoc: q_28_ = 3.815, p = 0.0111) animals.

### Viral expression of tau^P301L^ augments OXT immunofluorescence in 18-mo PVH males in the absence of neurodegeneration

Finally, we characterized whether tau^P301L^ expression dysregulated the PVH OXT system (**Fig. 4**). We were unable to compare across age groups as tissue was processed separately and therefore subject to batch effects. Consistent with a lack of tau^P301L^ expression in the PVH in hippocampocortical animals, there were no effects of such pathological expression on proportion of OXT neurons that were phosphorylated tau-positive, number of OXT neurons, or OXT immunofluorescence (**Fig. 4C,D,G,J**). In PVH animals, like hippocampocortical animals, only a small proportion of OXT neurons stained positive for phosphorylated tau (<35%; **Fig. 4C,E,F**), consistent with a previous report using pan-neuronal promoters in this brain

**Fig. 4:**
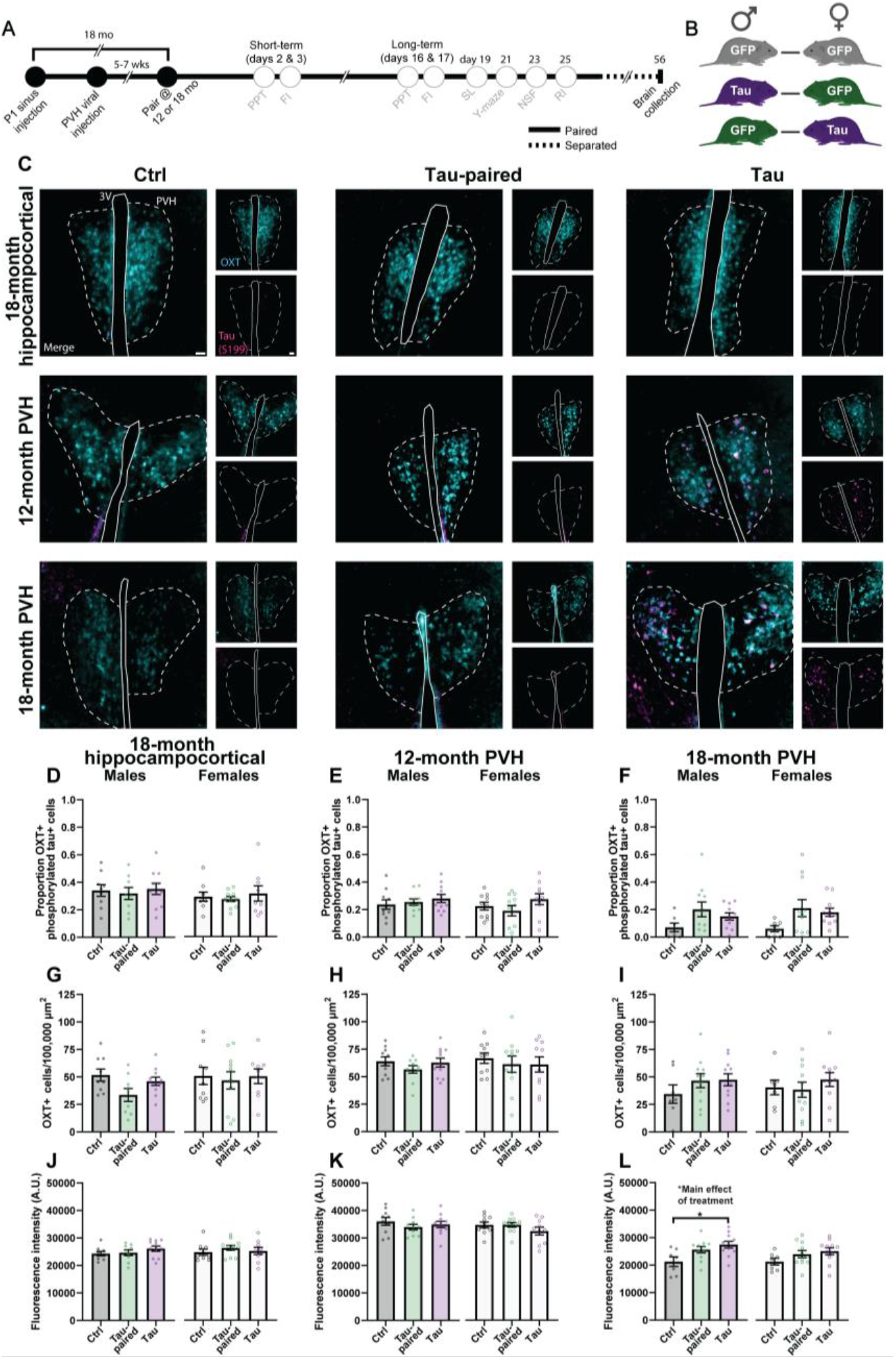
Tau^P301L^ expression increased OXT immunofluorescence in 18-month PVH males. **A** Experimental timeline. PVH: paraventricular hypothalamus. **B** Pairing scheme. **C** Representative images of OXT (cyan) and phosphorylated tau (S199, magenta) staining in the PVH from hippocampocortical and PVH targeted animals across the three pairing types (ctrl, tau-paired, and tau). Scale = 50 µm. **D-F** A majority of OXT cells remained free from phosphorylated tau across treatment groups. **G-I** There were no effects of tau^P301L^ expression on the number of PVH OXT neurons. **J-L** While 18-month hippocampocortical and 12-month PVH animals displayed no difference in OXT immunofluorescence, 18-month PVH males expressing tau^P301L^ pathology displayed an increase in OXT immunofluorescence compared to controls. 18-month PVH females displayed no change in OXT immunofluorescence. *p<0.05.

region^67^. Subsequent examinations revealed no differences in OXT neuron number or immunofluorescence in 12-month animals (**Fig. 4C,H,K**). Similarly, while there were no detectable differences in OXT cell number in 18-month PVH animals (**Fig. 4C,I**), OXT fluorescence intensity was affected in 18-month PVH males (One-way ANOVA, main effect of treatment: F_2,26_ = 4.895, p = 0.0157; **Fig. 4C,L**) where 18-month males expressing tau^P301L^ displayed elevated OXT immunofluorescence levels compared to controls (Tukey’s post-hoc: q_26_ = 4.395, p = 0.0122).

## Discussion

Despite reports that social withdrawal and other socially-relevant phenotypes are among the earliest symptoms of neurodegenerative disease^1,3^, there has been very little work focused on the mechanisms that drive these deficits and how they affect social relationships. This is particularly relevant given that strong social relationships facilitate health and well-being, and can slow disease progression^17,19–22^. Here, using monogamous prairie voles, we provide direct evidence that dementia-related tau^P301L^ disrupts pair bonding in an sex-, age-, and region-specific manner, establishing a model that will be crucial for understanding the complex interplay between social behavior, neurodegenerative disease, and hypothalamic function.

We found a striking regional dissociation in the effects of tau^P301L^ on bonding: hypothalamic, but not hippocampocortical, expression produced deficits in bonding behavior, despite the much shorter duration of PVH expression (∼5-7 weeks) compared to lifelong expression in hippocampocortical animals, which we necessarily transduced on postnatal day 1 to achieve widespread expression^65^. Across three tests that examine different aspects of bond-related behavior, deficits were primarily evident in social interaction and affiliation rather than aggression. Aging typically leads to faster bonding in male voles^54,55^, yet 18-month PVH tau^P301L^ males failed to form bonds even after long-term cohabitation. At a group level, these males exhibited weaker partner preference, driven primarily by a greater amount of time spent huddling with a stranger vole, which may suggest a loss of typical selectivity and/or discrimination that characterizes pair bonds. These males also exhibited reduced total huddle time, indicating an overall reduction in social interest and/or social drive. This was similarly reflected in reduced total social interaction with a partner even when animals were allowed to freely interact in the dyadic interaction test, although deficits in this test were also evident in 12-month male and female pairs containing tau^P301L^ animals. This suggests that dyadic interaction phenotypes may be more sensitive than partner preference and aligns with prior reports that the dyadic interaction test is partly dissociable from partner preference^54,66^. The convergence of social dysfunction in 18-month PVH tau^P301L^ males across tasks suggests that pathology produces pervasive social bonding disruptions that generalize across behavioral contexts. Finally, no significant differences in aggression were observed in the resident intruder test, suggesting PVH-mediated social dysfunction is relatively specific to affiliative behaviors. Together, these findings indicate that female bonding and overall sociability are comparatively resilient to tau^P301L^, and that the PVH is a major contributor to age-related, tau-induced attachment deficits in dementia primarily in males.

PVH tau^P301L^-induced deficits appear largely restricted to the social domain. We did not observe differences in arousal, anxiety-like behavior, or cognition. This is somewhat surprising given the multifaceted nature of the PVH, which is responsible for producing several neuropeptides that are important for stress responsivity (e.g. corticotropin releasing hormone and vasopressin) and cognition (oxytocin and vasopressin)^32,34–38,68,69^. It was equally surprising that hippocampocortical tau^P301L^ failed to yield detectable behavioral phenotypes. This raises the possibility that these circuits may be more resilient to tau^P301L^ expression than the PVH or that compensatory mechanisms may mitigate the functional consequences of such pathology over time. Voles are an outbred species unlike common laboratory rodents, and other broad genetic perturbations have similarly failed to produce robust changes in behavior even though they have strong effects in mice^70,71^. There may therefore be an element of genetic buffering to the effects of tau expression, but our data would suggest this occurs in an sex-, age-, and brain-region-dependent manner.

We also observed vicarious effects of tau^P301L^ expression predominately in males, establishing a model for asking how pathology in one partner affects the behavior of the other partner. 18-month males paired with PVH tau^P301L^ females took longer to form bonds, and their eventual bonds were weaker than those of controls. This occurred even though the tau^P301L^ females paired to these males didn’t exhibit deficits in social behavior that were detectable in the tests we employed. In contrast, female partners of tau^P301L^ males did not exhibit this vicarious disruption of social bonding, exhibiting partner preference despite the impaired social behavior of their partner. This finding may be particularly relevant for understanding the elevated caregiver burden reported among spouses^8–15^, and suggests that at least part of this burden may stem from changes in the relationship itself, rather than solely from the practical demands of caregiving^72^. Further development of this prairie vole model could be useful in defining the neural mechanisms of burden imposed by having a spouse with dementia.

The behavioral patterns we detected in voles parallel observations in human dementia patients, where men often display more numerous and severe changes in personality and behavioral symptoms than women^73–75^, while women exhibit greater behavioral resilience for equivalent neuropathology^76^. In addition, this pattern of sex differences may also reflect well-established differences in pair bonding in voles. Male voles typically take longer to form bonds and display lower levels of affiliative behavior than females^54,66^, suggesting that male bonds are potentially less robust and more susceptible to perturbations. This is consistent with prior work in which at least some types of stressors specifically disrupt male bonds^77,78^.

To gain a more refined understanding of the potential role of OXT in our observed social behavior deficits, we asked how our viral manipulations altered OXT neurons in tau^P301L^-expressing voles. Phosphorylated tau was detected primarily in non-OXT neurons, consistent with prior reports that the CMV promoter primarily drives transgene expression in non-OXT neurons^67^, though whether this extends to other peptidergic populations is unclear and may help explain the lack of behavioral effects outside the social domain. Despite the lack of phosphorylated tau in most OXT neurons, we observed increased OXT immunoreactivity in 18-month PVH male animals. Considering that OXT neuron number was not different, this finding suggests that tau expression may affect per-cell OXT content, reflecting an upregulation of OXT production. This is at odds with other rodent findings, which show a downregulation of OXT in dementia models^50,79–81^, but potentially clarifies human findings. The human PVH develops tau pathology during normal aging and across a variety of dementias^39–42^. Reports of outright cell loss and hypertrophy of the PVH in both aging and dementia are mixed^40,43–46^. Similar mixed findings have been reported for OXT levels in cerebrospinal fluid, serum, and brain samples from dementia patients^82–84^. Our data may therefore help reconcile these seemingly conflicting human findings: if OXT follows a biphasic or stage-dependent trajectory of dysfunction, then discrepancies across human studies could be due to differences in disease stage or pathology burden at time of sampling. The OXT upregulation we observe could be an early compensatory response that fails as the disease progresses, similar to that observed in other neuromodulatory subcortical systems^32,85^.

These results also suggest that PVH OXT circuitry may be less vulnerable to tau-induced dysfunction in females, which may help explain their preserved bonding capabilities despite equivalent phosphorylated tau expression. Previous studies directly comparing sexes found that females tend to show greater OXT neuron number and innervation, whereas males tend to have higher downstream OXT receptor expression^32,86,87^. Such elevated baseline OXT function in females may therefore buffer against tau-induced circuit dysfunction. Varying the length of viral expression or the age of testing might uncover pair bonding deficits in females, paralleling how resilience in humans tends to wane with advancing disease stage^88^. Alternatively, this reserve may not reflect resilience at the level of the PVH, but may reflect engagement of alternative circuits^89,90^.

There are several limitations to consider for the present work. Our behavioral assessments relied on well-established behavioral tasks, which may miss more nuanced behavioral deficits. Accordingly, applying pose-estimation and behavioral classification to unconstrained dyadic interaction could help reveal earlier and more nuanced changes in bonding. We also focused on how existing pathology affects bond formation. While fundamentally important, introducing pathology into voles with established bonds is needed to understand how bond maintenance and partner loss are affected by PVH tau. This may be particularly revealing for females, whose bonds tend to erode quicker than males following partner separation^54^. Also, while our results hint at changes in male OXT production upon PVH tau^P301L^ expression, more work is needed to determine parameters of OXT function that are dysregulated during disease, including measuring OXT cell activity and/or OXT release. Regardless, any OXT-based interventions will likely need to account disease subtype, stage, and sex, mirroring considerations that are increasingly being applied to therapeutics that target other neuromodulatory subcortical systems^31,32,91^. Finally, the hypothalamus is vulnerable to different pathologies based on dementia subtype^41^, and expressing these varied pathologies in the PVH may help differentiate their contributions to attachment deficits across different diseases.

Together, these findings establish that dementia-related tau pathology disrupts pair bonding in a manner that is sex-, age-, and region-specific, with PVH pathology selectively impairing bond formation in aged males, while leaving female bonding, and non-social behavior more broadly, largely unaffected. These results position the PVH, and potentially OXT circuity, as a key node linking dementia pathology to social dysfunction, while also revealing that healthy partners are behaviorally responsive to an affected partner. This underscores that pair bonds, and supporting neural circuits, are dynamic, bidirectional units vulnerable to disease on both sides of a relationship. Prairie voles thus offer a tractable, clinically relevant model for dissecting the neural mechanisms linking social bonds to dementia progression, with direct implications for understanding, and eventually treating, the social burdens of disease.

## Acknowledgements

This manuscript is dedicated to my parents, Marc and Christine Kelberman, and sister, Marissa Kelberman. My father passed away on August 29, 2026, after a battle with cancer, and my mother was his primary caregiver over the past few years. Their unwavering support has allowed me to pursue my dreams and become the scientist I am today, and I hope I have made them proud. May this manuscript serve as a reminder to cherish the moments we share with the people who matter most. I am optimistic that my work, along with the work of others in this space, will help build a future where we get to enjoy a few more of these moments with our loved ones.

We would like to thank Jessica Abazaris and the Office of Animal Resources (OAR) staff at the University of Colorado Boulder for their excellent care of our vole colony. This work was funded by Simons Foundation International SFI-AN-NC-AB-Fellow-00023830 (M.A.K.), the National Institutes of Health U01 NS131406-S1 (K.E.W.), T32 GM133289-01A1 (K.E.W.), F31 MH144897-0 (K.E.W.), U24 AG072701 NEW Brain Aging 2023 Pilot Program (Z.R.D.), and R01 MH125423 (Z.R.D.), the University of Colorado Boulder Undergraduate Research Opportunities Program (R.E.I. and E.G.A.), and the Renée Crown Wellness Institute Undergraduate Research Fellowship (R.E.I.).

This paper was typeset with the bioRxiv word template by @Chrelli: www.github.com/chrelli/bioRxiv-word-template.

## Author contributions

M.A.K. conceived and performed experiments, analyzed data, and wrote/revised the manuscript. K.E.W. assisted with behavioral experiments, optimized immunohistochemistry staining, collected images for analysis, and provided edits for the manuscript. R.E.I. assisted with behavioral experiments and analyzed immunohistochemistry data.

E.G.A. optimized immunohistochemistry staining and analysis methods.

Y.L.W. assisted with behavioral experiments and provided general colony management. M.P. assisted with behavioral experiments and provided edits for the manuscript. Z.R.D. conceived experiments, provided all project resources, and wrote/revised the manuscript.

## Materials and Methods

### Animals

Prairie voles were bred in-house at the University of Colorado Boulder in a colony originating from wild caught animals from Illinois.

Animals were housed in same-sex groups of 2-4 in static rodent cages until being paired. Animals had free access to water and rabbit chow supplemented with sunflower seeds with cotton nestlets, igloos, and tubes for enrichment. Housing conditions were maintained at 23 – 26 °C on a 14:10 hr light:dark cycle. All procedures were performed during the light cycle and approved by the Institutional Animal Care and Use committee at the University of Colorado Boulder.

### Stereotactic Surgery

For neonate injections, P1 pups were placed in a plastic cup with nesting material from the home cage on a heating pad. Pups were placed in the index finger of a nitrile glove and placed into a plastic weigh boat filled with ice^65^. When the absence of movement to light skin pinch was detected, the pups’ head was placed into a modified blunt ear bars on a stereotax. The incision site was disinfected with 3 rounds of alternating iodine and alcohol and a small incision was made in the skin above each transverse sinus. A 26G x 1/2’ sterile hypodermic needle was used to make a small hole directly above each sinus and a glass pipette was lowered through the skull into the sinus. Animals were injected with AAV_php.eB_-CMV-GFP or AAV_php.eB_-CMV-GFP-tau^P301L^ (2 µL/hemisphere at 10 nL/s). All viral vectors were produced by the Canadian Optogenetics and Vectorology Foundry Viral Vector Core, and the tau^P301L^ construct was based on Addgene plasmid 46908^92^. The skin was closed using vetbond and topical lidocaine was applied to the incision site. Animals were placed on a heating pad in a weigh boat with home cage nesting material until movement was detected and pink coloring was observed, after which they were returned to their mothers and observed for full recovery. Animals were monitored for the following 72 hrs.

For PVH injections, 5-7 weeks prior to pairing with an opposite-sex partner, animals were injected with AAV_1_-CMV-GFP or AAV_1_- CMV-GFP-tau^P301L^. Voles were anesthetized with isoflurane (4% induction with 1.5-3% maintenance) at an oxygen flow rate of 1 L/min. Voles were head-fixed in a stereotaxic frame (Kopf instruments) and ophthalmic ointment was used to lubricate the eyes. Body temperature was maintained using a heating pad and anesthesia depth was monitored by breathing rate and toe/tail pinch response. Fur was removed with a shaver, and the incision site was disinfected with 3 rounds of alternating iodine and alcohol. An incision was made along the midline and the skull was levelled. A single drill hole was made along the midline above the approximate location of the PVH a glass pipette attached to a nanoject was used to bilaterally deliver virus (AP: -0.7 ML: ±0.1 DV: -5.75 from bregma; 200 nL/hemisphere at 2 nL/s). After each injection, the glass pipette was left in place for 10 min to allow the virus to diffuse. The incision site was sutured and secured with vetbond, along with application of topical lidocaine and antibiotic ointment. Animals were provided fluid support (1 mL warmed saline S.C.) and an analgesic (4 mg/kg SR meloxicam S.C.) and placed on a heating pad while monitoring recovery. Animals were observed for 72 hrs and provided supplementary support, as needed.

In addition to viral injections, females were tubally ligated, either 3-4 weeks prior to pairing in neonate injected animals or at the time of viral injection in adults to avoid pregnancy confounds while leaving hormonal function intact. Fallopian tubes were cauterized and bisected via lumbar incisions on each side of the body. Body cavities were closed with absorbable sutures, and the skin was closed with staples after which topical lidocaine and antibiotic ointment were applied. Staples were removed 10-14 days after surgery.

### Experimental timeline

A full experimental timeline and pairing scheme can be found in **Fig. 1A,B**. Experiments were performed on opposite-sex, non-sibling pairs at 12- or 18-months in PVH injected animals and on 18-month hippocampocortical animals. Pairs consisted of male GFP animals paired with female GFP animals (control), male GFP animals paired with female tau^P301L^ animals, and male tau^P301L^ animals paired with female GFP animals (see **Table 1** for details on experimental groups). Animals were paired (day 0) and partner preference tests were performed on both members of the pair following 2 days (short-term) and 16 days (long-term/2 wks) of cohabitation. Animals were tested on a free interaction test on days 3 and 17. After completing free interaction on day 17, animals were tested on sleep latency (day 19), Y-maze (day 21), novelty suppressed feeding (day 23), and resident intruder tests (day 25). Animals were then separated (day 26) and brains were harvested and analyzed for tau and oxytocin immunofluorescence 30 days later (day 56).

### Behavioral procedures

*Partner preference tests:* Testing was performed as described previously^54,66,93^. Partner and stranger non-sibling animals were briefly anesthetized with isoflurane to tether them to opposite ends of a three- chamber plexiglass arena (76.0 cm l, 20.0 cm w, 30 cm t). Stranger voles were reproductively intact, sexually naïve, and housed in same-sex cages of 2-4 animals. Tethers were composed of chains with zip ties on either end, with one end connected around the animal’s neck and the other placed through a hole in the chamber wall to secure the animal in place, allowing all animals access to food and water throughout the 3 hr test. Experimental animals were placed into the center chamber that was separated from the other chambers at the beginning of the test with opaque dividers. Opaque dividers were removed and the experimental animal was allowed to freely explore the arena for 3 hrs. Animals were tracked via an overhead Panasonic WVCP304 camera or Logitech C925e webcam. Behavior was scored using TopScan High-Throughput software v3.0 (Cleversys Inc.) following previously established protocols^54,66,94^. Both voles within a pair were tested in partner preference tests back-to-back on each test day, separated by approximately 1 hr. Test order (male or female) and side the partner was tethered to were randomized and counterbalanced within an age and test group. The main outcome variables analyzed from these tests were total time spent huddling with the partner and stranger animal, and partner preference (partner huddle time/[partner huddle time + stranger huddle time).

*Free interaction:* The day after each partner preference test, animals were recorded during unconstrained dyadic partner interactions in a plexiglass arena, as described previously ^54,66^. Pairs were placed on opposite sides of the two-chamber arena (50.7 cm l, 20.0 cm w, and 30.0 cm t) separated by an opaque divider. The divider was removed, and animals were allowed to freely explore the chamber for 3 hrs which was recorded using overhead Logitech C925e webcams. Social contact between animals was scored using TopScan High-Throughput software v3.0 (Cleversys Inc.) using methods optimized within our lab^54,66^. We measured time spent in close social contact (defined by setting the “joint motion” parameter to <5).

*Sleep latency:* Animals were placed in a familiar chamber (50.7 cm l, 20.0 cm w, 30.0 cm t) 2-3 hrs after lights on after gentle handling in clear plastic cups and recorded for 6 hrs. Sleep latency was scored as 2 min of continuous sleep posture followed by sleep in 75% of the next 8 min, as previously described^95^. Sleep latency is an EEG validated measure of sleep in rodents^95,96^.

*Y-maze:* Animals were placed in the center of an opaque acrylic Y-maze (35 cm arm l, 20 cm arm h, 5 cm arm w) and allowed to freely explore for 8 min while being video recorded. The sequence and total number of arm entries were recorded. Percent alternation was calculated as: (number of consecutive arm entries into three different arms)/(total arm entries minus two).

*Novelty suppressed feeding:* Animals were food deprived in a clean cage for 2-3 hrs and then placed into an open field (40 cm x 40 cm x 40 cm) with a circular dish (25 cm diameter) filled with 2 cm of water and a food pellet atop an inverted 100 mL beaker placed in the center of the dish. Notably, these conditions have been optimized to detect increased/decreased latency to eat in voles, deviating slightly from those previously employed in mice and rats which yielded a floor effect^28,97^. Latency for voles to bite the foot pellet was recorded with a 20 min time out. Immediately after, animals were returned to their home cage with a pre-weighed food pellet and the latency to eat and amount eaten within 30 min were recorded.

*Resident intruder:* Same sex intruders that were approximately age- and size-matched were placed in the residents’ home cage (11 in x 8 in x 6.5 in) and behavior was video recorded for 10 min, until there was 30 s of total fighting, or 5 bouts of tumble fighting, whichever came first.

### Immunohistochemistry, imaging, and cell counts

At the end of behavioral testing, animals were injected with a mixture of ketamine/xylazine and perfused with 1x PBS followed by 4% paraformaldehyde. Brains were stored in 4% paraformaldehyde overnight and then transferred to 30% sucrose. Brains were sliced at 40 µm on a Leica SM2010R microtome and slices were stored in 0.01% sodium azide in 1x PBS at 4°C until staining.

For staining, 1-3 sections per animal encompassing the PVH were washed 3 × 5 min in 1x PBS and then incubated for 1 hr at room temperature in blocking solution (0.2% Triton-X, 10% normal goat serum in 1x PBS). Sections were then incubated in 1:500 rabbit anti-S199 (ThermoFisher Scientific, 44-734G) primary antibody in blocking buffer for 48 hrs. After 48 hrs, sections were washed 3 x 5 min in 1x PBS and then incubated in 1:500 Alexa Fluor™ goat anti-rabbit 568 secondary antibody (ThermoFisher Scientific, A-11036) for 2 hrs at room temperature. Sections were again washed 3 x 5 min in 1x PBS and then incubated in 10% normal rabbit serum in 0.2% Triton-X in 1x PBS for 2 hrs at room temperature. Sections were then washed 3 x 5 min in 1x PBS and then incubated in AffiniPure® Fab Fragment Goat Anti-Rabbit (Jackson ImmunoResearch Laboratories Inc., 111-007-003) for 2 hrs at room temperature. Sections were washed 3 x 5 min in 1x PBS and then incubated for 24 hrs in 1:500 chicken anti-GFP (Aves Labs, GFP-1020) and rabbit anti-OXT (Immunostar, 20068) primary antibodies diluted in blocking buffer. The next day, sections were washed 3 x 5 min in 1x PBS and then incubated in 1:500 Alexa Fluor™ goat anti-chicken 488 (abcam, ab150169) and goat anti-rabbit 405 (ThermoFisher Scientific, A48254) secondary antibodies for 2 hrs at room temperature. Sections underwent a final round of 3 x 5 min washes in 1x PBS at which point they were mounted on Diamond White Glass microscope slides, coverslipped with ProLong™ Diamond Antifade Mountant (ThermoFisher Scientific, P36970), and sealed with nail polish.

Images were acquired on an Olympus iX83 wide field slide scanner with a Chroma Multi LED filter set (Cat: 69401) at 20x for quantification with consistent exposures times (DAPI 150 ms, FITC 50 ms, TRITC 50 ms). Images used for quantification were imported into ImageJ/Fiji (v1.54s). For OXT cell counts, a rolling ball subtraction of 50 was used to reduce image background. Images were grayscaled and thresholded using the Otsu method (0–50). The watershed function was applied to separate individual cells and analyze particles (size = 50- infinity, circularity = 0.3-1.00) was performed to count the number of OXT cells within a manually drawn region of interest (ROI) that encompassed the PVH. The manually drawn ROI was also used to measure fluorescence intensity of the raw OXT image. OXT cell counts per area were averaged across all sections for an animal. Outlines of the OXT cells were saved as a separate ROI and applied to the raw tau fluorescence channel. The total number of phosphorylated tau inclusions within the PVH and the number of OXT cells that were also positive for phosphorylated tau were manually counted by two blinded individuals and averaged per slice and then per animal.

### Statistical analyses

All data are presented as mean ± SEM and behavioral data were scored by observers blinded to experimental group. Because our primary interest was the effects of tau pathology on pair bonding, rather than potential sex differences in bonding, we examined sexes separately. Molecular data was analyzed with a one-way ANOVA if it was normally distributed (Shapiro-Wilk test) and standard deviations did not differ according to a Brown-Forsythe test. A Tukey’s multiple comparisons test was used for pairwise comparisons. We used a Kruskal-Wallis test to analyze non-normally distributed data, with a Dunn’s multiple comparison test for pairwise comparisons. If only standard deviations were not equivalent between groups, a Brown- Forsythe ANOVA was used. Partner preference and free interaction tests also included repeated measures for the short- and long-term timepoints. Analyses were separated into individual groups (12-month PVH, 18-month PVH, and 18-month hippocampocortical) meaning that treatment refers to the virus and pair type that an animal was a part of (GFP-GFP/control, GFP paired to tau^P301L^, or tau^P301L^ paired to GFP). Timepoint refers to the specific test day that animals were tested on (2 days/short-term and 2 wks/long-term). When there were interactions between treatment and timepoint, we followed up with Holm-Šidák post- hoc tests with two families (timepoint and treatment) for correction. All statistics were performed and graphs were made in GraphPad Prism (version 10.6.1), with figures being constructed in Adobe Illustrator 2025 (version 29.8.9). Figures 1B, 2B, 3B-H, and 4B, as well as Supplemental Figures 2B and 3B, were created in Biorender. Donaldson, Z. (2026) https://BioRender.com/shduqhd. All statistical analyses are available in **Table S2**.

## Data availability

All data associated with this paper has been deposited on and is publicly available through Zenodo: https://zenodo.org/records/22097178?preview=1&token=eyJhbGciOiJIUzUxMiJ9.eyJpZCI6IjNhMGI4YjQ3LTJmMWItNGRjYy1iNWFmLWNmMWE3MzRkZGQ4ZiIsImRhdGEiOnt9LCJyYW5kb20iOiIzMDk0YjYwM2EyMzUxMjdhNWI3YTEwOGYwY2E4MTc5ZSJ9.NzWC3U_9yzE-o46yUhJjZoLq9WunD8iWc1WH-1y-hPhnxtQVp6hhsA7bDyIQkDrbfbLWujeXUkF_aVTcRBLEFw

---

**Fig. S1:**
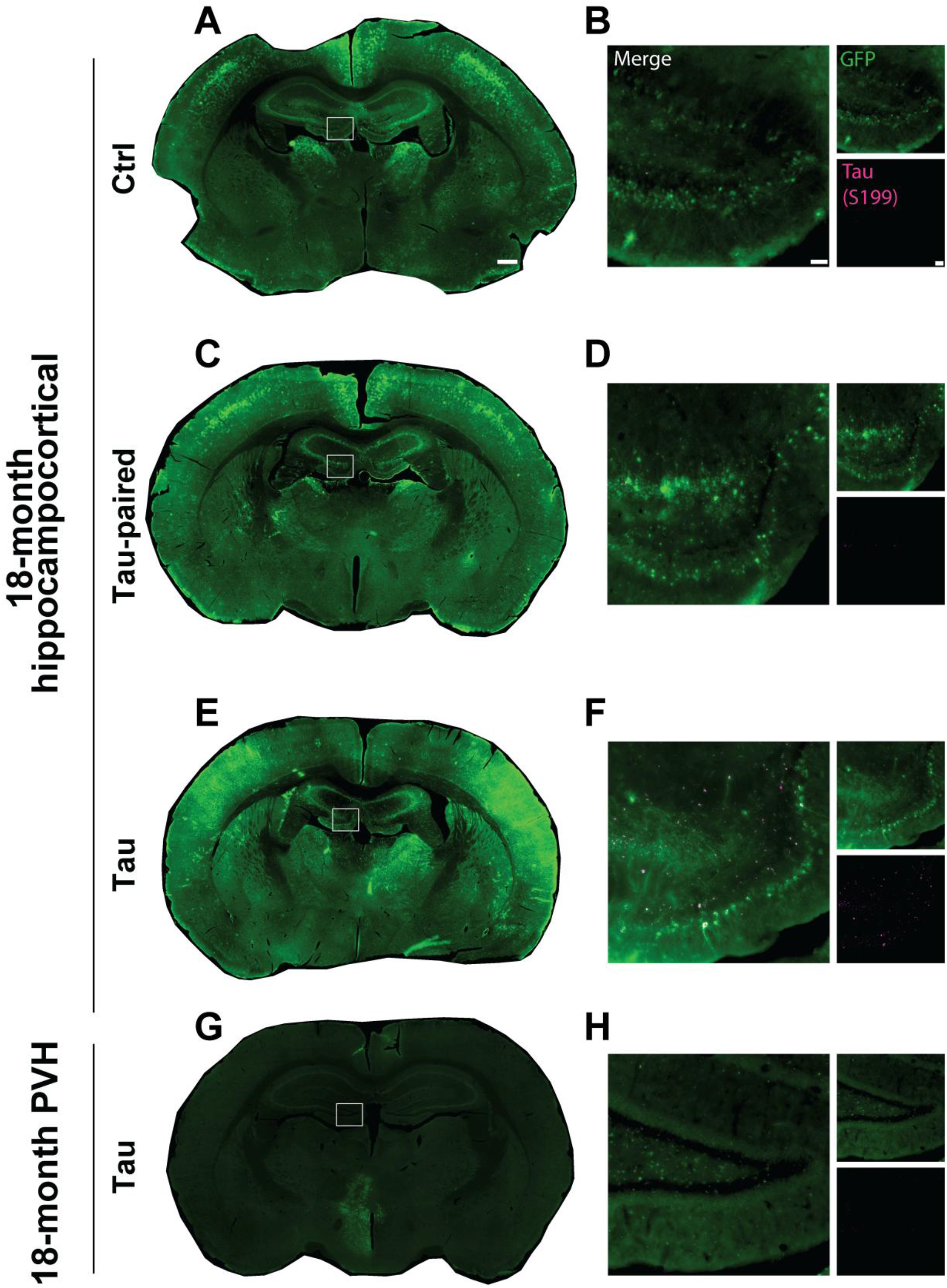
Neonate sinus viral injections result in broad hippocampocortical expression while PVH viral injections result in focal expression. **A-F** Representative images of whole hippocampal slices and zoom-ins of the dentate gyrus from control (A,B), tau-paired (C,D), and tau^P301L^ (E,F) animals. **G-H** Representative images of a whole hippocampal slice and a zoom-in of the dentate gyrus from an 18-month PVH tau^P301L^ animal. Scale = 500 µm (whole slice) and 50 µm (inset). Hippocampal slices display GFP (green) and zoom-ins display both GFP (green) and phosphorylated tau (S199, magenta).

**Fig. S2:**
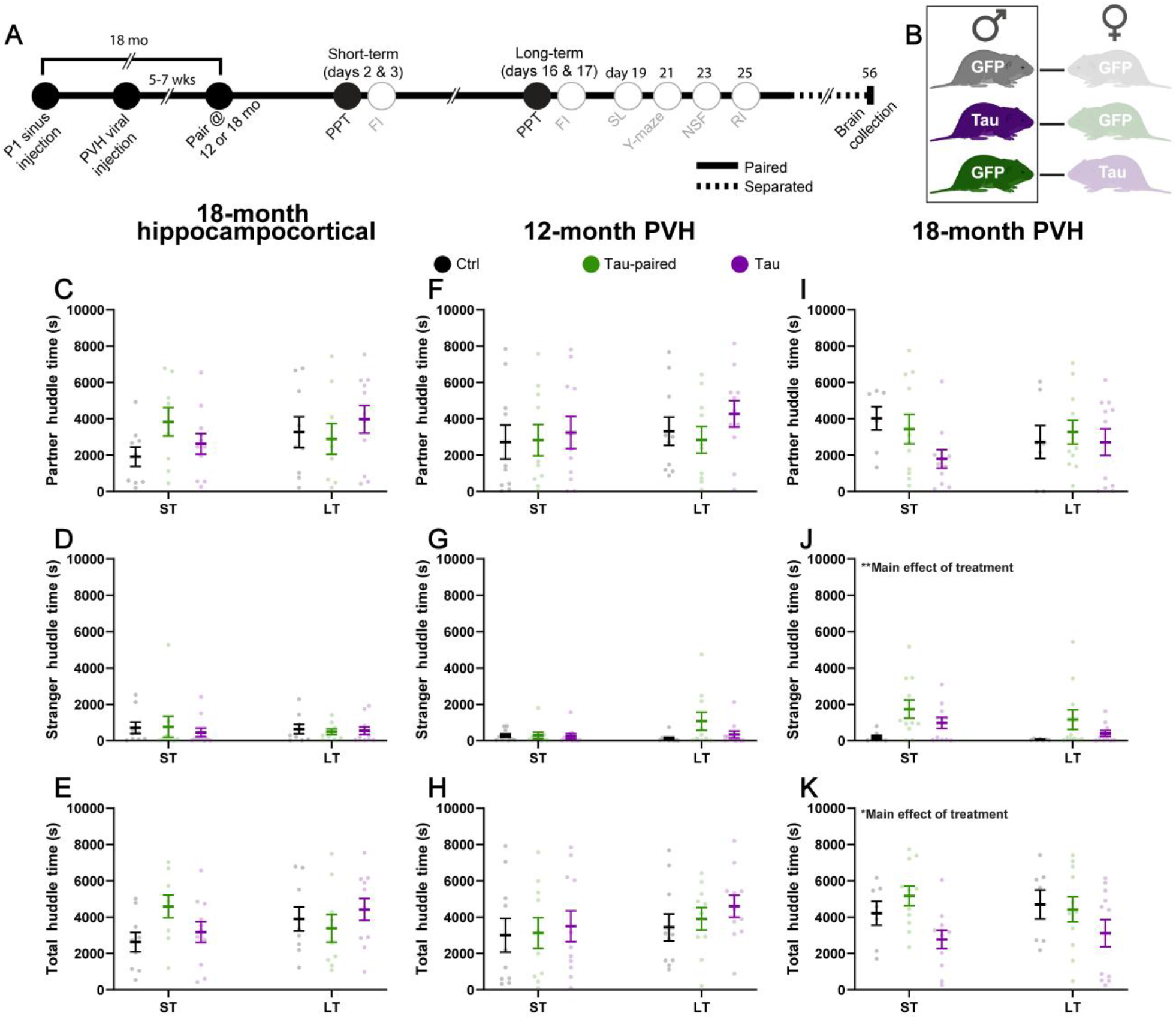
Huddle times during partner preference tests reveal additional deficits in bonding in 18-month PVH males. **A** Experimental timeline. PVH: paraventricular hypothalamus, PPT: partner preference test. **B** Pairing scheme. **C-E** Partner (C), stranger (D), and total huddle times (E) from 18-month hippocampocortical males. There were no differences in any of these metrics. **F-H** Partner (F), stranger (G), and total huddle times (H) from 12-month PVH males. There were no differences in any of these metrics. **I-K** Partner (I), stranger (J), and total huddle times (K) from 18-month PVH males. 18-month PVH tau-paired and tau^P301L^ males displayed elevated stranger huddle time, and tau^P301L^ males displayed the lowest amount of total huddle time. n = 7-11 voles per group. *p<0.05, **p<0.01.

**Fig. S3:**
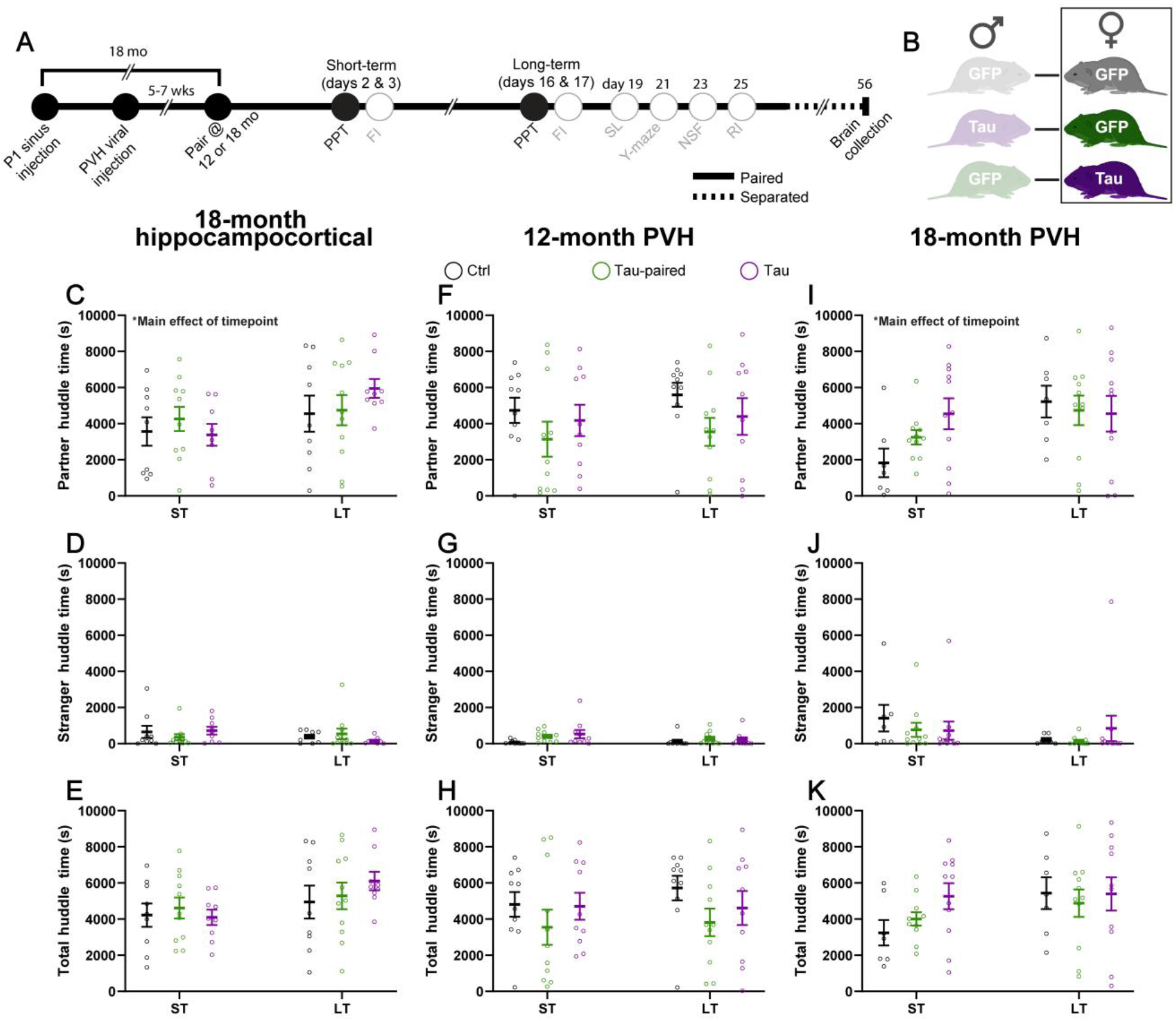
Huddle times during partner preference tests reveal additional effects of timepoint on bonding in females. **A** Experimental timeline. PVH: paraventricular hypothalamus, PPT: partner preference test. **B** Pairing scheme. **C-E** Partner (C), stranger (D), and total huddle times (E) from 18-month hippocampocortical females. Females across groups tended to increase time spent huddling with a partner after long-term cohabitation. **F-H** Partner (F), stranger (G), and total huddle times (H) from 12-month PVH females. There were no differences in any of these metrics. **I-K** Partner (I), stranger (J), and total huddle times (K) from 18-month PVH females. Females across groups tended to increase time spent huddling with a partner after long-term cohabitation. n = 7-11 voles per group. *p<0.05.

## References

1. Ismail, Z. et al. Neuropsychiatric symptoms as early manifestations of emergent dementia: Provisional diagnostic criteria for mild behavioral impairment. Alzheimer’s & Dementia 12, 195–202 (2016).

2. Mendez, M. F. et al. Observation of Social Behavior in Frontotemporal Dementia. Am J Alzheimers Dis Other Demen 29, 215–221 (2014).

3. Jost, B. C. & Grossberg, G. T. The evolution of psychiatric symptoms in Alzheimer’s disease: a natural history study. J Am Geriatr Soc 44, 1078–1081 (1996).

4. Kar, N. Behavioral and psychological symptoms of dementia and their management. Indian J Psychiatry 51, S77–S86 (2009).

5. García-Martín, V., de Hoyos-Alonso, M. C., Delgado-Puebla, R., Ariza-Cardiel, G. & del Cura-González, I. Burden in caregivers of primary care patients with dementia: influence of neuropsychiatric symptoms according to disease stage (NeDEM project). BMC Geriatr 23, 525 (2023).

6. Creese, B. et al. Mild Behavioral Impairment as a Marker of Cognitive Decline in Cognitively Normal Older Adults. Am J Geriatr Psychiatry 27, 823–834 (2019).

7. Lyketsos, C. G. et al. Neuropsychiatric symptoms in Alzheimer’s disease. Alzheimers Dement 7, 532–539 (2011).

8. Harris, M. L., Titler, M. G. & Hoffman, G. J. Associations Between Alzheimer’s Disease and Related Dementias and Depressive Symptoms of Partner Caregivers. J Appl Gerontol 40, 772–780 (2021).

9. Sallim, A. B., Sayampanathan, A. A., Cuttilan, A. & Ho, R. Prevalence of Mental Health Disorders Among Caregivers of Patients With Alzheimer Disease. J Am Med Dir Assoc 16, 1034–1041 (2015).

10. Lou, Q. et al. Comprehensive analysis of patient and caregiver predictors for caregiver burden, anxiety and depression in Alzheimer’s disease. J Clin Nurs 24, 2668–2678 (2015).

11. Norton, M. C. et al. Increased Risk of Dementia When Spouse Has Dementia? The Cache County Study. J Am Geriatr Soc 58, 895–900 (2010).

12. Wu-Chung, E. L., Leal, S. L., Denny, B. T., Cheng, S. L. & Fagundes, C. P. Spousal caregiving, widowhood, and cognition: A systematic review and a biopsychosocial framework for understanding the relationship between interpersonal losses and dementia risk in older adulthood. Neurosci Biobehav Rev 134, 104487 (2022).

13. Choi, H. et al. Family Care Availability And Implications For Informal And Formal Care Used By Adults With Dementia In The US. Health Affairs 40, 1359–1367 (2021).

14. Reckrey, J. M., Bollens-Lund, E., Husain, M., Ornstein, K. A. & Kelley, A. S. Family Caregiving for Those With and Without Dementia in the Last 10 Years of Life. JAMA Intern Med 181, 278–279 (2021).

15. Alzheimer’s Association. Alzheimer’s Disease Facts and Figures. Alzheimer’s & https://doi.org/10.1002/alz.71345 doi:10.1002/alz.71345. Dementia (2026)

16. Salinas, J. et al. Association of Social Support With Brain Volume and Cognition. JAMA Netw Open 4, e2121122 (2021).

17. Costa-Cordella, S., Arevalo-Romero, C., Parada, F. J. & Rossi, A. Social Support and Cognition: A Systematic Review. Front. Psychol. 12, (2021).

18. Bennett, D. A., Schneider, J. A., Tang, Y., Arnold, S. E. & Wilson, R. S. The effect of social networks on the relation between Alzheimer’s disease pathology and level of cognitive function in old people: a longitudinal cohort study. Lancet Neurol 5, 406–412 (2006).

19. Sommerlad, A., Ruegger, J., Singh-Manoux, A., Lewis, G. & Livingston, G. Marriage and risk of dementia: systematic review and meta-analysis of observational studies. J Neurol Neurosurg Psychiatry 89, 231–238 (2018).

20. Gerritsen, L. et al. Influence of negative life events and widowhood on risk for dementia. Am J Geriatr Psychiatry 25, 766–778 (2017).

21. Luchetti, M. et al. A Meta-analysis of Loneliness and Risk of Dementia using Longitudinal Data from >600,000 Individuals. Nat Ment Health 2, 1350–1361 (2024).

22. Desai, R., John, A., Stott, J. & Charlesworth, G. Living alone and risk of dementia: A systematic review and meta-analysis. Ageing Research Reviews 62, 101122 (2020).

23. Tissot, C. et al. Association between regional tau pathology and neuropsychiatric symptoms in aging and dementia due to Alzheimer’s disease. Alzheimers Dement (N Y) 7, e12154 (2021).

24. Ismail, Z. et al. Affective and emotional dysregulation as pre-dementia risk markers: exploring the mild behavioral impairment symptoms of depression, anxiety, irritability, and euphoria. International Psychogeriatrics 30, 185–196 (2018).

25. Thal, D. R., Rüb, U., Orantes, M. & Braak, H. Phases of Aβ-deposition in the human brain and its relevance for the development of AD. Neurology 58, 1791–1800 (2002).

26. Braak, H., Braak, E. & Bohl, J. Staging of Alzheimer-Related Cortical Destruction. Eur Neurol 33, 403–408 (2008).

27. Braak, H., Thal, D. R., Ghebremedhin, E. & Del Tredici, K. Stages of the Pathologic Process in Alzheimer Disease: Age Categories From 1 to 100 Years. J Neuropathol Exp Neurol 70, 960–969 (2011).

28. Kelberman, M. A. et al. Consequences of Hyperphosphorylated Tau in the Locus Coeruleus on Behavior and Cognition in a Rat Model of Alzheimer’s Disease. Journal of Alzheimer’s Disease 86, 1037–1059 (2022).

29. Khan, K. M. et al. Human tau-overexpressing mice recapitulate brainstem involvement and neuropsychiatric features of early Alzheimer’s disease. Acta Neuropathol Commun 11, 57 (2023).

30. Vercruysse, P., Vieau, D., Blum, D., Petersén, Å. & Dupuis, L. Hypothalamic Alterations in Neurodegenerative Diseases and Their Relation to Abnormal Energy Metabolism. Front Mol Neurosci 11, 2 (2018).

31. Ehrenberg, A. J. et al. Priorities for research on neuromodulatory subcortical systems in Alzheimer’s disease: Position paper from the NSS PIA of ISTAART. Alzheimers Dement 19, 2182–2196 (2023).

32. Rae, R. J. et al. Sex differences in neuromodulatory subcortical systems and their implications for Alzheimer’s disease. Alzheimer’s & Dementia 22, e71291 (2026).

33. Khan, N. S. et al. Behavioral and Neurophysiological Implications of Pathological Human Tau Expression in Serotonin Neurons. ACS Chem. Neurosci. 15, 932–943 (2024).

34. Donaldson, Z. R. & Young, L. J. Oxytocin, Vasopressin, and the Neurogenetics of Sociality. Science 322, 900–904 (2008).

35. Bosch, O. J., Nair, H. P., Ahern, T. H., Neumann, I. D. & Young, L. J. The CRF System Mediates Increased Passive Stress-Coping Behavior Following the Loss of a Bonded Partner in a Monogamous Rodent. Neuropsychopharmacology : official publication of the American College of Neuropsychopharmacology 34, 1406 (2009).

36. Bosch, O. J. et al. Oxytocin in the nucleus accumbens shell reverses CRFR2-evoked passive stress-coping after partner loss in monogamous male prairie voles. Psychoneuroendocrinology 64, 66–78 (2016).

37. Cho, M., C, D. a., R, W. j. & S, C. c. The effects of oxytocin and vasopressin on partner preferences in male and female prairie voles (Microtus ochrogaster). Behav. Neurosci. 113, 1071–1079 (1999).

38. Young, L. J. & Wang, Z. The neurobiology of pair bonding. Nat Neurosci 7, 1048–1054 (2004).

39. Diodati, D., Cyn-Ang, L., Kertesz, A. & Finger, E. Pathologic evaluation of the supraoptic and paraventricular nuclei in dementia. Can J Neurol Sci 39, 213–219 (2012).

40. Son, G. et al. Spatially conserved pathoprotein profiling in the human suprachiasmatic nucleus in progressive Alzheimer disease stages. 2024.03.07.584000 Preprint at 10.1101/2024.03.07.584000 (2024).

41. Stewart, C. A. & Finger, E. C. Chapter 7 - The supraoptic and paraventricular nuclei in healthy aging and neurodegeneration. in Handbook of Clinical Neurology (eds Swaab, D. F., Kreier, F., Lucassen, P. J., Salehi, A. & Buijs, R. M.) vol. 180 105–123 (Elsevier, 2021).

42. Swaab, D. F. et al. Tau and ubiquitin in the human hypothalamus in aging and Alzheimer’s disease. Brain Res 590, 239–249 (1992).

43. de Lacalle, S., Iraizoz, I. & Gonzalo, L. M. Cell loss in supraoptic and paraventricular nucleus in Alzheimer’s disease. Brain Research 609, 154–158 (1993).

44. Fliers, E., Swaab, D. F., W. Pool Chr. & Verwer, R. W. H. The vasopressin and oxytocin neurons in the human supraoptic and paraventricular nucleus; changes with aging and in senile dementia. Brain Research 342, 45–53 (1985).

45. Goudsmit, E., Hofman, M. A., Fliers, E. & Swaab, D. F. The supraoptic and paraventricular nuclei of the human hypothalamus in relation to sex, age and Alzheimer’s disease. Neurobiol Aging 11, 529–536 (1990).

46. Vogels, O. J., Broere, C. A. & Nieuwenhuys, R. Neuronal hypertrophy in the human supraoptic and paraventricular nucleus in aging and Alzheimer’s disease. Neurosci Lett 109, 62–67 (1990).

47. El-Ganainy, S. O. et al. Intranasal Oxytocin Attenuates Cognitive Impairment, β-Amyloid Burden and Tau Deposition in Female Rats with Alzheimer’s Disease: Interplay of ERK1/2/GSK3β/Caspase-3. Neurochem Res 47, 2345–2356 (2022).

48. Takahashi, J. et al. Oxytocin reverses Aβ-induced impairment of hippocampal synaptic plasticity in mice. Biochem Biophys Res Commun 528, 174–178 (2020).

49. Takahashi, J. et al. Intracerebroventricular administration of oxytocin and intranasal administration of the oxytocin derivative improve β-amyloid peptide (25–35)-induced memory impairment in mice. Neuropsychopharmacology Reports 42, 492–501 (2022).

50. Selles, M. C. et al. Oxytocin attenuates microglial activation and restores social and non-social memory in APP/PS1 Alzheimer model mice. iScience 26, (2023).

51. Edwards, H. M. et al. Sex-dependent effects of acute stress on amyloid-β in male and female mice. Brain 146, 2268–2274 (2023).

52. Li, C.-Y. et al. Effect of Endogenous Arginine-Vasopressin Arising from the Paraventricular Nucleus on Learning and Memory Functions in Vascular Dementia Model Rats. BioMed Research International 2017, 3214918 (2017).

53. Carter, C. S. & Getz, L. L. Monogamy and the prairie vole. Sci. Am 268, 100–106 (1993).

54. Kelberman, M. A., Winther, K. E., Medvedeva, Y. M. & Donaldson, Z. R. Aging leads to sex-dependent effects on pair bonding and increased number of oxytocin-producing neurons in monogamous prairie voles. Hormones and Behavior 166, 105647 (2024).

55. Kenkel, W. M., Perkeybile, A. M., Yee, J. R. & Carter, C. S. Rewritable fidelity: How repeated pairings and age influence subsequent pair-bond formation in male prairie voles. Hormones and Behavior 113, 47–54 (2019).

56. Powell, J. M., Garvin, M. M., Lee, N. S. & Kelly, A. M. Behavioral trajectories of aging prairie voles (Microtus ochrogaster): Adapting behavior to social context wanes with advanced age. PLoS ONE 17, e0276897 (2022).

57. Grippo, A. J. et al. Behavioral and neuroendocrine consequences of disrupting a long-term monogamous social bond in aging prairie voles. Stress 24, 239–250 (2021).

58. Schaepe, C. A. & Hiura, L. C. Behavioral sex differences in the developmental emergence and selectivity of partner preferences in prairie voles. Horm Behav 173, 105778 (2025).

59. Luong, G., Charles, S. T. & Fingerman, K. L. Better with age: Social relationships across adulthood. Journal of Social and Personal Relationships 28, 9–23 (2011).

60. Charles, S. T. & Piazza, J. R. Memories of social interactions: Age differences in emotional intensity. Psychology and Aging 22, 300–309 (2007).

61. Houlden, H. et al. Frequency of tau mutations in three series of non-Alzheimer’s degenerative dementia. Ann Neurol 46, 243–248 (1999).

62. Poorkaj, P. et al. Frequency of Tau Gene Mutations in Familial and Sporadic Cases of Non-Alzheimer Dementia. Arch Neurol 58, 383–387 (2001).

63. Dumanchin, C. et al. Segregation of a missense mutation in the microtubule-associated protein tau gene with familial frontotemporal dementia and parkinsonism. Hum Mol Genet 7, 1825–1829 (1998).

64. Hutton, M. et al. Association of missense and 5’-splice-site mutations in tau with the inherited dementia FTDP-17. Nature 393, 702–705 (1998).

65. Hamodi, A. S., Martinez Sabino, A., Fitzgerald, N. D., Moschou, D. & Crair, M. C. Transverse sinus injections drive robust whole-brain expression of transgenes. eLife 9, e53639 (2020).

66. Brusman, L. E. et al. Emergent intra-pair sex differences and organized behavior in pair bonded prairie voles (Microtus ochrogaster). Genes Brain Behav 21, e12786 (2022).

67. Rajamani, K. T. et al. Efficiency of cell-type specific and generic promoters in transducing oxytocin neurons and monitoring their neural activity during lactation. Scientific Reports 11, (2021).

68. Jiang, Z. & Tong, Q. Hypothalamic CRH neurons: A crossroad between stress and metabolism. Current Opinion in Endocrine and Metabolic Research 26, 100384 (2022).

69. Smith, A. S. & Wang, Z. Hypothalamic oxytocin mediates social buffering of the stress response. Biol Psychiatry 76, 281–288 (2014).

70. Berendzen, K. M. et al. Oxytocin receptor is not required for social attachment in prairie voles. Neuron 111, 787–796.e4 (2023).

71. Pobbe, R. L. H. et al. Oxytocin receptor knockout mice display deficits in the expression of autism-related behaviors. Horm Behav 61, 436–444 (2012).

72. Monin, J. K., Levy, B., Doyle, M., Schulz, R. & Kershaw, T. The impact of both spousal caregivers’ and care recipients’ health on relationship satisfaction in the Caregiver Health Effects Study. J Health Psychol 24, 1744–1755 (2019).

73. Silvestri, C. et al. Sex Differences in the Severity and Progression of Neuropsychiatric Symptoms Across Different Dementia Types. Neurol Clin Pract 14, e200299 (2024).

74. Eikelboom, W. S. et al. Sex differences in neuropsychiatric symptoms in Alzheimer’s disease dementia: a meta-analysis. Alzheimer’s Research & Therapy 14, 48 (2022).

75. Pengo, M. et al. Sex influences clinical phenotype in frontotemporal dementia. Neurol Sci 43, 5281–5287 (2022).

76. Illán-Gala, I. et al. Sex differences in the behavioral variant of frontotemporal dementia: A new window to executive and behavioral reserve. Alzheimers Dement 17, 1329–1341 (2021).

77. Jones, C. E. et al. Early-life sleep disruption increases parvalbumin in primary somatosensory cortex and impairs social bonding in prairie voles. Sci Adv 5, eaav5188 (2019).

78. DeVries, A. C., DeVries, M. B., Taymans, S. E. & Carter, C. S. The effects of stress on social preferences are sexually dimorphic in prairie voles. Proceedings of the National Academy of Sciences 93, 11980–11984 (1996).

79. Usmani, S. S. et al. Targeting the hypothalamus for modeling age-related DNA methylation and developing OXT-GnRH combinational therapy against Alzheimer’s disease-like pathologies in male mouse model. Nat Commun 15, 9419 (2024).

80. Jackson, H. M., Soto, I., Graham, L. C., Carter, G. W. & Howell, G. R. Clustering of transcriptional profiles identifies changes to insulin signaling as an early event in a mouse model of Alzheimer’s disease. BMC Genomics 14, 831 (2013).

81. Li, J. et al. Oxytocin Intervention Mitigates Pathological and Behavioral Impairments in APP/PS1 Mice Subjected to Early Social Isolation. CNS Neurosci Ther 31, e70511 (2025).

82. Raskind, M. A. et al. Cerebrospinal fluid vasopressin, oxytocin, somatostatin, and beta-endorphin in Alzheimer’s disease. Arch Gen Psychiatry 43, 382–388 (1986).

83. Mazurek, M. F., Beal, M. F., Bird, E. D. & Martin, J. B. Oxytocin in Alzheimer’s disease: postmortem brain levels. Neurology 37, 1001–1003 (1987).

84. Petekkaya, E., Burakgazi, G., Kuş, B., Melek, İ. M. & Arpacı, A. Comparative study of the volume of the temporal lobe sections and neuropeptide effect in Alzheimer’s patients and healthy persons. Int J Neurosci 131, 725–734 (2021).

85. Kelberman, M. A. et al. Age-dependent dysregulation of locus coeruleus firing in a transgenic rat model of Alzheimer’s disease. Neurobiology of Aging 125, 98–108 (2023).

86. Dumais, K. M. & Veenema, A. H. Vasopressin and oxytocin receptor systems in the brain: sex differences and sex-specific regulation of social behavior. Front Neuroendocrinol 40, 1–23 (2016).

87. Lu, Q. & Hu, S. Chapter 5 - Sex differences of oxytocin and vasopressin in social behaviors. in Handbook of Clinical Neurology (eds Swaab, D. F., Kreier, F., Lucassen, P. J., Salehi, A. & Buijs, R. M.) vol. 180 65–88 (Elsevier, 2021).

88. Arenaza-Urquijo, E. M. et al. Sex and gender differences in cognitive resilience to aging and Alzheimer’s disease. Alzheimer’s & Dementia 20, 5695–5719 (2024).

89. de Vries, G. J. Sex differences in vasopressin and oxytocin innervation of the brain. in Advances in Vasopressin and Oxytocin — From Genes to Behaviour to Disease (eds Neumann, I. D. & Landgraf, R.) vol. 170 17–27 (Elsevier, 2008).

90. Shvareva, V. A., Berendzen, K. M., Bender, K. J. & Manoli, D. S. Dimorphic substrates, monomorphic outcomes: Social behavior neural circuit tuning in social monogamy. Progress in Neurobiology 264, 102948 (2026).

91. Weinshenker, D. Long Road to Ruin: Noradrenergic Dysfunction in Neurodegenerative Disease. Trends in Neurosciences 41, 211–223 (2018).

92. Hoover, B. R. et al. Tau mislocalization to dendritic spines mediates synaptic dysfunction independently of neurodegeneration. Neuron 68, 1067–1081 (2010).

93. Sadino, J. M. et al. Prolonged partner separation erodes nucleus accumbens transcriptional signatures of pair bonding in male prairie voles. eLife 12, e80517 (2023).

94. Ahern, T. H., Modi, M. E., Burkett, J. P. & Young, L. J. Evaluation of two automated metrics for analyzing partner preference tests. Journal of neuroscience methods 182, 180–8 (2009).

95. Porter-Stransky, K. A. et al. Noradrenergic transmission at alpha1-adrenergic receptors in the ventral periaqueductal gray modulates arousal. Biol Psychiatry 85, 237–247 (2019).

96. Hunsley, M. S. & Palmiter, R. D. Norepinephrine-Deficient Mice Exhibit Normal Sleep-Wake States but have Shorter Sleep Latency after Mild Stress and Low Doses of Amphetamine. Sleep 26, 521–526 (2003).

97. Lustberg, D. et al. Noradrenergic circuits in the forebrain control affective responses to novelty. Psychopharmacology 237, 3337–3355 (2020).

