## Supplemental Table 1 for "Hypothalamic tau^P301L^ pathology disrupts pair bonding in monogamous prairie voles"

| **Age** | **Experimental Group** | **N** | **Mean age at pairing (wks; M : F)** | **Range age at pairing (wks; M : F)** |
| --- | --- | --- | --- | --- |
| 18-month neonate | GFP-GFP | 9 | 74.9 : 76.8 | 74 – 76 : 74 – 80 |
|  | Male GFP-female tau^P301L^ | 9 | 74.6 : 78.9 | 67 – 83 : 71 – 82 |
|  | Male tau^P301L^-female GFP | 11 | 77.4 : 78.5 | 74 – 80 : 74 – 81 |
| 12-month PVH | GFP-GFP | 10 | 51.3 : 50.1 | 47 – 55 : 41 – 56 |
|  | Male GFP-female tau^P301L^ | 10 | 52.8 : 51.0 | 50 – 56 : 45 – 56 |
|  | Male tau^P301L^-female GFP | 11 | 51.1 : 50.7 | 49 – 54 : 45 – 58 |
| 18-month PVH | GFP-GFP | 7 | 80.0 : 79.9 | 77 – 82 : 77 – 82 |
|  | Male GFP-female tau^P301L^ | 11 | 80.1 : 80.0 | 73 – 84 : 74 – 85 |
|  | Male tau^P301L^-female GFP | 11 | 79.8 : 80.1 | 73 – 87 : 72 – 86 |

**Table S1. Demographics of experimental groups**
